# Phylogenetic analysis supports a link between DUF1220 domain number and primate brain expansion

**DOI:** 10.1101/018077

**Authors:** Fabian Zimmer, Stephen H. Montgomery

## Abstract

The expansion of DUF1220 domain copy number during human evolution is a dramatic example of rapid and repeated domain duplication. However, the phenotypic relevance of DUF1220 dosage is unknown. Although patterns of expression, homology and disease associations suggest a role in cortical development, this hypothesis has not been robustly tested using phylogenetic methods. Here, we estimate DUF1220 domain counts across 12 primate genomes using a nucleotide Hidden Markov Model. We then test a series of hypotheses designed to examine the potential evolutionary significance of DUF1220 copy number expansion. Our results suggest a robust association with brain size, and more specifically neocortex volume. In contradiction to previous hypotheses we find a strong association with postnatal brain development, but not with prenatal brain development. Our results provide further evidence of a conserved association between specific loci and brain size across primates, suggesting human brain evolution occurred through a continuation of existing processes.

## Introduction

The molecular targets of selection favoring brain expansion during human evolution have been sought by identifying dramatic, lineage-specific shifts in evolutionary rate. The increase in DUF1220 domains during human evolution provides one of the most dramatic increases in copy number (Popesco et al., 2006; Dumas et al., 2012). A single copy of this protein domain is found in *PDE4DIP* in most mammalian genomes. In primates, this ancestral domain has been duplicated many times over, reaching its peak abundance in humans where several hundred DUF1220 domains exist across 20-30 genes in the Nuclear Blastoma Breakpoint Family (NBPF) (Vandepoele et al., 2005; Dumas et al., 2012). The majority of these map to 1q21.1, a chromosomal region with complex, and unstable genomic architecture (O’Bleness et al., 2012, 2014).

Interspecific DUF1220 counts show a pattern of phylogenetic decay with increasing distance from humans (Popesco et al., 2006; Dumas and Sikela, 2009; Dumas et al., 2012). In humans, DUF1220 dosage has also been linked to head circumference (Dumas et al., 2012), and severe neurodevelopmental disorders, including autism spectrum disorders (ASD) and microcephaly (Dumas et al., 2012; Davis et al., 2014). The severity of ASD impairments is also correlated with 1q21.1 DUF1220 copy number suggesting a dosage effect (Davis et al., 2014). Taken together, these observations led to the suggestion that the expansion of DUF1220 copy number played a primary role in human brain evolution (Dumas and Sikela, 2009; Keeney et al., 2014a).

The strength of this hypothesis is difficult to assess given the paucity of information on the developmental function of NBPF genes and the DUF1220 domain. DUF1220 domains are highly expressed during periods of cortical neurogenesis, suggesting a potential role in prolonging the proliferation of neural progenitors by regulating centriole and microtubule dynamics to control key cell fate switches critical for neurogenesis (Keeney et al., 2014b). *PDE4DIP*, which contains the ancestral DUF1220 domain, does indeed associate with the spindle poles (Popesco et al., 2006) and is homologous to *CDK5RAP2*, a centrosomal protein essential for neural proliferation (Bond et al., 2005; Buchman et al., 2010), which co-evolved with brain mass across primates (Montgomery et al., 2011).

Two previous analyses reported a significant association between DUF1220 copy number and brain mass, cortical neuron number (Dumas et al., 2012), cortical gray and white matter, surface area and gyrification (Keeney et al., 2014b). However, several limitations in these analyses restrict confidence in the results. First, DUF1220 copy number was assessed across species using a BLAT/BLAST analysis with a query sequence from humans, which introduces a bias that may partly explain the observed phylogenetic decay. Second, counts were not restricted to those domains occurring in functional exonic sequence. Finally, the analyses were limited to a small number of species (4-8 primates), and did not correct for phylogenetic non-independence (Felsenstein, 1985) or autocorrelation between traits.

Here, we use nucleotide Hidden Markov Models (HMMER3; Eddy, 2011) to more accurately query the DUF1220 domain number of distantly related genomes. After filtering these counts to limit the analysis to exonic sequence, we use phylogenetic comparative methods that correct for non-independence to test whether DUF1220 copy number is robustly associated with brain size, whether this is due to an association with pre- or postnatal brain development, and whether the association is specific to the neocortex.

## Results

We find evidence that CM-associated exonic DUF1220 counts (Table 1) are associated with brain mass across primates (n = 12, posterior mean =1.927, 95% CI = 0.800-3.040, p_MCMC_ = 0.001). This association is robust to the exclusion of *Homo* (posterior mean =1.271, 95% CI = 0.490-2.019, p_MCMC_ = 0.003), and found when hominoids (n = 5, posterior mean = 3.679, 95% CI = 0.966-6.258, p_MCMC_ = 0.018) or anthropoids (n = 9, posterior mean = 2.019, 95% CI = 0.352-3.684, p_MCMC_ = 0.010) are analyzed alone, suggesting a consistent phylogenetic association. When body mass is included as a co-factor in the model, the positive association is restricted to brain mass (Table 2a).

**Table 1:** DUF1220 count data.

| Species | O'Bleness et al.<br>(2012) | nHMM |  |
| --- | --- | --- | --- |
|  |  | whole<br>genome | functional exonic<br>with CM promoter |
| *Homo sapiens* | 272 | 302 | 262 |
| *Pan troglodytes* | 125 | 138 | 34 |
| *Gorilla gorilla* | 99 | 97 | 32 |
| *Pongo abelii* | 92 | 101 | 27 |
| *Nomascus leucogenys* | 53 | 59 | 6 |
| *Papio anubis* | - | 75 | 15 |
| *Chlorocebus sabaeus* | - | 48 | 16 |
| *Macaca mulatta* | 35 | 74 | 10 |
| *Callithrix jacchus* | 31 | 75 | 9 |
| *Tarsius syrichta* | - | 47 | 2 |
| *Microcebus murinus* | 2 | 4 | 1 |
| *Otolemur garnettii* | 3 | 4 | 1 |

**Table 2:** MCMCglmm results of multivariate models.

**a) Brain mass and body mass**
| **Model** | **Posterior mean** | **95% CI** | **p<sub>MCMC</sub>** |
| --- | --- | --- | --- |
| 1. log(brain mass) | 4.105 | 2.163 - 6.000 | 0.001 |
| + log(body mass) | -1.986 | -3.544 - -3.900 | 0.988 |

**b) Prenatal and postnatal growth**
| **Model** | **Posterior mean** | **95% CI** | **p<sub>MCMC</sub>** |
| --- | --- | --- | --- |
| 1. log(prenatal brain growth) | -2.158 | -4.471 - 0.106 | 0.967 |
| + log(postnatal brain growth) | 3.319 | 1.470 - 4.982 | 0.002 |
| 2. log(postnatal brain growth) | 2.910 | 1.641 - 4.151 | <0.001 |
| + log(postnatal body growth) | -1.241 | -2.442 - -0.052 | 0.977 |

**c) Brain regions**
| **Model** | **Posterior mean** | **95% CI** | **p<sub>MCMC</sub>** |
| --- | --- | --- | --- |
| 1. log(neocortex volume) | 5.961 | 0.720 - 11.173 | 0.014 |
| + log(RoB volume) | -5.817 | -13.322 - 1.120 | 0.953 |
| 2. log(cerebellum volume) | 3.699 | -5.857 - 12.611 | 0.186 |
| + log(RoB volume) | -2.435 | -13.869 - 10.132 | 0.681 |
| 3. log(neocortex volume) | 6.076 | -0.139 - 12.5712 | 0.025 |
| + log(cerebellum volume) | -0.369 | -9.5128 - 8.961 | 0.526 |
| + log(RoB volume) | -5.494 | -15.814 - 5.288 | 0.872 |

Separation of pre- and postnatal development specifically links DUF12220 number to postnatal brain growth. Analysed separately, the association with prenatal brain growth is weaker (n = 11, posterior mean =1.758, 95% CI = −0.039-3.543, p_MCMC_ = 0.023) than with postnatal brain growth (posterior mean =1.839, 95% CI = 0.895-2.808, p_MCMC_ = 0.001). If both traits are included in the same model, only the positive association with postnatal brain growth remains (Table 2b). Multiple regression analysis also confirms the association is specific to postnatal brain growth, rather than postnatal body growth (Table 2b).

Finally, we examined the hypothesized relationship with neocortex volume (e.g. Keeny et al., 2014a,b), but also consider cerebellum volume, as this region co-evolves with the neocortex (Barton and Harvey, 2000), has expanded in apes (Barton and Venditti, 2014), and shows high levels of NBPF expression (Popesco et al., 2006). When the rest-of-the-brain (RoB) is included as a co-factor, to account for variation in overall brain size, a positive association is found for neocortex volume but not cerebellum volume (Table 2c).

## Discussion

Our phylogenetic analyses support the hypothesis that the increase in DUF1220 number co-evolves with brain mass, and contributes to the proximate basis of primate brain evolution. We also find evidence of specific associations with neocortex volume and postnatal brain growth. Previous hypotheses concerning the phenotypic relevance of DUF1220 domain number have focused on their possible contribution to neurogenesis (Dumas and Sikela, 2009; Keeney et al., 2014a; b). This is supported by homology to genes with known functions in cell cycle dynamics (Popesco et al., 2006; Thornton and Woods, 2009), relevant spatial and temporal expression patterns (Keeney et al., 2014b), and an effect on the proliferation of neuroblastoma cell cultures (Vandepoele et al., 2008). However, a direct effect of variation in DUF1220 domain number on neural proliferation has not been demonstrated (Keeney et al., 2015).

If DUF1220 domains do regulate neurogenesis, we would expect them to coevolve with prenatal brain growth, as cortical neurogenesis is restricted to prenatal development (Bhardwaj et al., 2006). Our results instead suggest a robust and specific relationship with postnatal brain development. Existing data on DUF1220 domain function suggest two potential roles that may explain this association: i) a contribution to axonogenesis via initiating and stabilizing microtubule growth in dendrites; and ii) a potential role in apoptosis during brain maturation. Both hypotheses are consistent with the reported association between variation in DUF1220 dosage and ASD (Davis et al., 2014). Indeed, an emphasis on postnatal brain growth is potentially more relevant for ASD, which develops postnatal, accompanied by a period of accelerated brain growth (Courchesne et al., 2001).

Microtubule assembly is essential for dendritic growth and axonogenesis (Conde and Cáceres, 2009). *PDE4DIP*, which contains the ancestral DUF1220 domain, has known functions in microtubule nucleation, growth, and cell migration (Roubin et al., 2013). There is also evidence NBPF1 interacts with a key regulator of Wnt signaling (Vandepoele et al., 2010), which has important roles in neuronal differentiation, dendritic growth and plasticity (Inestrosa and Varela-Nallar, 2014). Consistent with this function, DUF1220 domains are highly expressed in the cell bodies and dendrites of adult neurons (Popesco et al., 2006). A role for DUF1220 domains in synaptogenesis could potentially explain the association with ASD severity (Davis et al., 2014). ASDs are associated with abnormalities in cortical minicolumns (Casanova et al., 2002) and cortical white matter (Hazlett et al., 2005; Courchesne et al., 2011), both of which suggest a disruption of normal neuronal maturation (Courchesne and Pierce, 2005; Minshew and Williams, 2007).

Alternatively, NBPF genes are also known to interact with NF-κB (Zhou et al., 2013), a transcription factor implicated in tumor progression, with a range of roles including apoptosis and inflammation (Karin and Lin, 2002; Perkins, 2012). Postnatal apoptosis has a significant influence on brain growth (Kuan et al., 2000; Polster et al., 2003; Madden et al., 2007), including regulating neuronal density (Sanno et al., 2010), and apoptotic genes may have been targeted by selection in relation to primate brain expansion (Vallender and Lahn, 2006). Disruption of apoptosis causes microcephaly (Poulton et al., 2011), potentially explaining the association between DUF1220 dosage and head circumference (Dumas et al., 2012). The association of NF-κB with inflammatory diseases (Tak et al., 2001) is also intriguing, given the growing evidence that the inflammatory response is linked to the risk and severity of ASD (Meyer et al., 2011; Depino, 2012).

If DUF1220 domain number does contribute to the evolution of postnatal brain growth, this contrasts with results of previously studied candidate genes with known roles in neurogenesis that co-evolve with prenatal brain growth (Montgomery et al., 2011). This suggests a two-component model of brain evolution where selection targets one set of genes to bring about an increase in neuron number (e.g. Montgomery et al., 2011; Montgomery and Mundy, 2012a;b), and an independent set of genes to optimize neurite growth and connectivity (e.g. Charrier et al., 2012). NBPF genes may fall into the latter category. This two-component model is consistent with comparative analyses that indicate pre- and postnatal brain development evolve independently, and must therefore be relatively free of reciprocal pleiotropic effects (Barton and Capellini, 2011).

Finally, these results add further evidence that many of the genetic changes that contribute to human evolution will be based on the continuation or exaggeration of conserved gene-phenotype associations that contribute to primate brain evolution (Montgomery et al., 2011; Scally et al., 2012). Understanding the commonalities between human and non-human primate brain evolution is therefore essential to understand the genetic differences that contribute the derived aspects of human evolution.

## Materials and methods

### Counting DUF1220 domains

HMMER3.1b (Eddy, 2011) was used to build a Hidden Markov Model (HMM) from the DUF1220 (PF06758) seed alignment stored in the PFAM database (Finn et al., 2014). The longest isoforms for all proteomes of 12 primate genomes from Ensembl v.78 (Cunningham et al., 2014) (Figure 1A), were searched using the protein DUF1220 HMM (hmmsearch, E-value < 1e-10) (Table S1). We extracted the corresponding cDNA regions to build a DUF1220 nucleotide profile HMM (nHMM), allowing for more sensitive analysis across a broad phylogenetic range. The DUF1220 nHMM was used to search the complete genomic DNA for all 12 species. These counts were filtered to remove any DUF1220 domains not located in annotated exonic sequence, or located in known pseudogenes.

**Figure 1:**
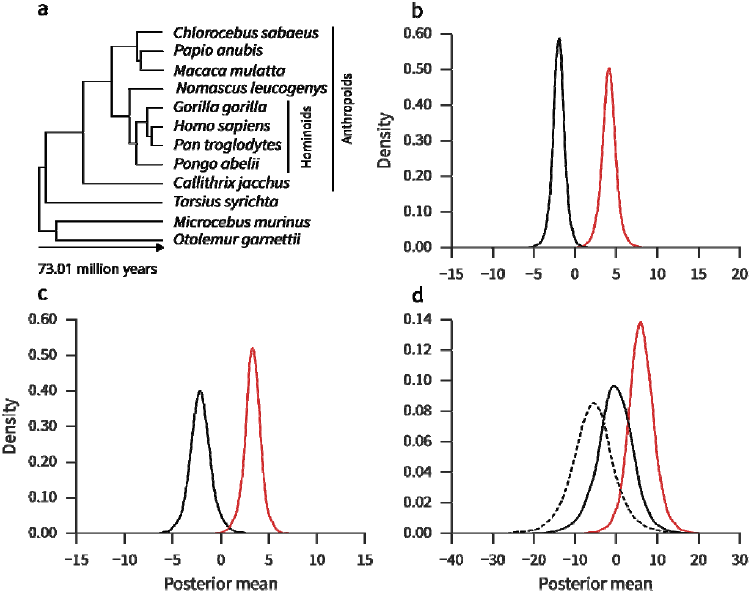
A) Phylogeny of Ensembl primates. B) Posterior means of the association between DUF1220 count and brain mass (red) and body mass (black). C) Posterior means of the association between DUF1220 count and postnatal brain growth (red) and prenatal brain growth (black). D) Posterior means of the association between DUF1220 count and neocortex volume (red), cerebellum volume (solid black) and rest-of-brain volume (dashed black).

We next filtered our counts to limit them to exonic sequence in close proximity to the NBPF-specific Conserved-Mammal (CM) promoter (O’Bleness et al. 2012). To do so, we built a nucleotide HMM for the CM promoter based on a MAFFT (Katoh et al., 2002) alignment of the 900bp CM region upstream of human genes NBPF4, NBPF6 and NBPF7. Using this CM promoter nHMM, we searched 1000bp up- and downstream of genes containing DUF1220 domains for significant CM promoter hits (nhmmer, E-value < 1e-10). This provided final counts for DUF1220 domains within exonic regions and associated with the CM promoter (Table 1). These counts were used in subsequent phylogenetic analyses. In the Supplementary Information we compare our counts with previous estimates and discuss possible sources of error. All scripts and data used in the analysis are freely available from: https://github.com/qfma/duf1220

## Phylogenetic gene-phenotype analysis

Phylogenetic multivariate generalized mixed models were implemented using a Bayesian approach in MCMCglmm (Hadfield, 2010), to test for phylogenetically-corrected associations between DUF1220 counts and *log*-transformed phenotypic data (Table S2). All analyses were performed using a Poisson distribution, as recommended for count data (O’Hara and Kotze, 2010), with uninformative, parameter expanded priors for the random effect (G: V = 1,n ν = 1, alpha.ν = 0, alpha.V = 1000; R: V = 1, ν = 0.002) and default priors for the fixed effects. Phylogenetic relationships were taken from the 10k Trees project (Arnold et al., 2010). We report the posterior mean of the co-factor included in each model and its 95% confidence intervals (CI), and the probability that the parameter value is >0 (p_MCMC_) as we specifically hypothesize a positive association (Dumas et al., 2012). Alternative data treatments lead to similar conclusions (Supplementary Information).

## Acknowledgments and funding information

We thank James Sikela, Majesta O’Bleness, Chris Venditti, Charlotte Montgomery, Andrew Moore and Jarrod Hadfield for advice and comments, and Judith Mank’s lab at UCL for support. SHM thanks the Leverhulme Trust for funding.

## References

Arnold C, Matthews LJ, Nunn CL. 2010. The 10kTrees website: A new online resource for primate phylogeny. Evol Anthropol 19:114–118.

Barton RA, Capellini I. 2011. Maternal investment, life histories, and the costs of brain growth in mammals. Proc Natl Acad Sci USA 108:6169–6174.

Barton RA, Harvey PH. 2000. Mosaic evolution of brain structure in mammals. Nature 405:1055–8.

Barton RA, Venditti C. 2014. Report Rapid Evolution of the cerebellum in humans and other great apes. Curr Biol 24:2440–2444.

Bhardwaj RD, Curtis MA, Spalding KL, Buchholz BA, Fink D, Björk-Eriksson T, Nordborg C, Gage FH, Druid H, Eriksson PS, Frisén J. 2006. Neocortical neurogenesis in humans is restricted to development. Proc Natl Acad Sci USA 103:12564–12568.

Bond J, Roberts E, Springell K, Lizarraga SB, Scott S, Higgins J, Hampshire DJ, Morrison EE, Leal GF, Silva EO, Costa SMR, Baralle D, Raponi M, Karbani G, Rashid Y, Jafri H, Bennett C, Corry P, Walsh CA, Woods CG. 2005. A centrosomal mechanism involving CDK5RAP2 and CENPJ controls brain size. Nat Genet 37:353–355.

Buchman JJ, Tseng HC, Zhou Y, Frank CL, Xie Z, Tsai LH. 2010. *CDK5RAP2* interacts with pericentrin to maintain the neural progenitor pool in the developing neocortex. Neuron 66:386–402.

Casanova MF, Buxhoeveden DP, Cohen M, Switala AE, Roy EL. 2002. Minicolumnar pathology in dyslexia. Ann Neurol 52:108–110.

Charrier C, Joshi K, Coutinho-Budd J, Kim JE, Lambert N, De Marchena J, Jin WL, Vanderhaeghen P, Ghosh A, Sassa T, Polleux F. 2012. Inhibition of *SRGAP2* function by its human-specific paralogs induces neoteny during spine maturation. Cell 149:923–935.

Conde C, Cáceres A. 2009. Microtubule assembly, organization and dynamics in axons and dendrites. Nat Rev Neurosci 10:319–332.

Courchesne E, Karns C, Davis H, Ziccardi R, Carper R, Tigue Z, Chisum HJ, Moses P, Pierce K, Lord C, Lincoln A, Pizzo S, Schreibman L, Haas R, Akshoomoff N, Courchesne R. 2011. Unusual brain growth patterns in early life in patients with autistic disorder: An MRI study. Neurology 76:2111.

Courchesne E, Pierce K. 2005. Why the frontal cortex in autism might be talking only to itself: Local over-connectivity but long-distance disconnection. Curr Opin Neurobiol 15:225–230.

Cunningham F, Amode MR, Barrell D, Beal K, Billis K, Brent S, Carvalho-Silva D, Clapham P, Coates G, Fitzgerald S, et al. 2014. Ensembl 2015. Nucleic Acids Res 43:D662–D669.

Davis JM, Searles VB, Anderson N, Keeney J, Dumas L, Sikela JM. 2014. DUF1220 Dosage is linearly associated with increasing severity of the three primary symptoms of autism. PLoS Genet 10:1–5.

Depino AM. 2012. Peripheral and central inflammation in autism spectrum disorders. Mol Cell Neurosci 53:69–76.

Dumas L, Sikela JM. 2009. DUF1220 domains, cognitive disease, and human brain evolution. Cold Spring Harb Symp Quant Biol 74:375–382.

Dumas LJ, O’Bleness MS, Davis JM, Dickens CM, Anderson N, Keeney JG, Jackson J, Sikela M, Raznahan A, Giedd J, Rapoport J, Nagamani SSC, Erez A, Brunetti-Pierri N, Sugalski R, Lupski JR, Fingerlin T, Cheung SW, Sikela JM. 2012. DUF1220-domain copy number implicated in human brain-size pathology and evolution. Am J Hum Genet 91:444–454.

Eddy SR. 2011. Accelerated profile HMM searches. PLoS Comput Biol 7.

Felsenstein J. 1985. Phylogenies and the Comparative Method. Am Nat 125:1–5.

Hadfield JD. 2010. MCMC methods for multi-response generalized linear mixed models: The MCMCglmm R package. J Stat Softw 33:1–22.

Hazlett HC, Poe M, Gerig G, Smith RG, Provenzale J, Ross A, Gilmore J, Piven J. 2005. Magnetic resonance imaging and head circumference study of brain size in autism: birth through age 2 years. Arch Gen Psychiatry 62:1366–1376.

Inestrosa NC, Varela-Nallar L. 2014. Wnt signalling in neuronal differentiation and development. Cell Tissue Res 359:215–223.

Karin M, Lin A. 2002. NF-κB at the crossroads of life and death. Nat Immunol 3:221–227.

Katoh K, Misawa K, Kuma K, Miyata T. 2002. MAFFT: a novel method for rapid multiple sequence alignment based on fast Fourier transform. Nucleic Acids Res 30:3059–3066.

Keeney J, Dumas L, Sikela J. 2014a. The case for DUF1220 domain dosage as a primary contributor to anthropoid brain axpansion. Front Hum Neurosci 8:1–11.

Keeney JG, Davis JM, Siegenthaler J, Post MD, Nielsen BS, Hopkins WD, Sikela JM. 2014b. DUF1220 protein domains drive proliferation in human neural stem cells and are associated with increased cortical volume in anthropoid primates. Brain Struct Funct 1–8.

Keeney JG, O’Bleness MS, Anderson N, Davis JM, Arevalo N, Busquet N, Chick W, Rozman J, Hölter SM, Garrett L, Horsch M, Beckers J, Wurst W, Klingenspor M, Restrepo D, de Angelis MH, Sikela JM. 2014c. Generation of mice lacking DUF1220protein domains: effects on fecundity and hyperactivity. Mamm Genome 26:33–42.

Kuan CY, Roth KA, Flavell RA, Rakic P. 2000. Mechanisms of programmed cell death in the developing brain. Trends Neurosci 23:291–297.

Madden SD, Donovan M, Cotter TG. 2007. Key apoptosis regulating proteins are down-regulated during postnatal tissue development. Int J Dev Biol 51:415–425.

Meyer U, Feldon J, Dammann O. 2011. Schizophrenia and autism: both shared and disorder-specific pathogenesis via perinatal inflammation? Pediatr Res 69:26–33.

Minshew NJ, Williams DL. 2007. The new neurobiology of autism. Arch Neurol 64:945–950.

Montgomery SH, Capellini I, Venditti C, Barton RA, Mundy NI. 2011. Adaptive evolution of four microcephaly genes and the evolution of brain size in anthropoid primates. Mol Biol Evol 28:625–638.

Montgomery SH, Mundy NI. 2012a. Evolution of *ASPM* is associated with both increases and decreases in brain size in primates. Evolution 66:927–32.

Montgomery SH, Mundy NI. 2012b. Positive selection on *NIN*, a gene involved in neurogenesis, and primate brain evolution. Genes, Brain Behav 11:903–910.

O’Bleness M, Searles VB, Dickens CM, Astling D, Albracht D, Mak ACY, Lai YYY, Lin C, Chu C, Graves T, Kwok P-Y, Wilson RK, Sikela JM. 2014. Finished sequence and assembly of the DUF1220-rich 1q21 region using a haploid human genome. BMC Genomics 15:387.

O’Bleness MS, Dickens CM, Dumas LJ, Kehrer-Sawatzki H, Wyckoff GJ, Sikela JM. 2012. Evolutionary history and genome organization of DUF1220 protein domains. G3 (Bethesda) 2:977–86.

O’Hara RB, Kotze DJ. 2010. Do not log-transform count data. Methods Ecol Evol 1:118–122.

Perkins ND. 2012. The diverse and complex roles of NF-κB subunits in cancer. Nat Rev Cancer 12(2): 121–132.

Polster BM, Robertson CL, Bucci CJ, Suzuki M, Fiskum G. 2003. Postnatal brain development and neural cell differentiation modulate mitochondrial Bax and BH3 peptide-induced cytochrome c release. Cell Death Differ 10:365–370.

Popesco MC, Maclaren EJ, Hopkins J, Dumas L, Cox M, Meltesen L, McGavran L, Wyckoff GJ, Sikela JM. 2006. Human lineage-specific amplification, selection, and neuronal expression of DUF1220 domains. Science 313:1304–1307.

Poulton CJ, Schot R, Kia SK, Jones M, Verheijen FW, Venselaar H, De Wit MCY, De Graaff E, Bertoli-Avella AM, Mancini GMS. 2011. Microcephaly with simplified gyration, epilepsy, and infantile diabetes linked to inappropriate apoptosis of neural progenitors. Am J Hum Genet 89:265–276.

Roubin R, Acquaviva C, Chevrier V, Sedjaï F, Zyss D, Birnbaum D, Rosnet O. 2013. Myomegalin is necessary for the formation of centrosomal and Golgi-derived microtubules. Biol Open 2:238–50.

Sanno H, Shen X, Kuru N, Bormuth I, Bobsin K, Gardner H a R, Komljenovic D, Tarabykin V, Erzurumlu RS, Tucker KL. 2010. Control of postnatal apoptosis in the neocortex by RhoA-subfamily GTPases determines neuronal density. J Neurosci 30:4221–4231.

Scally A, Dutheil JY, Hillier LW, Jordan GE, Goodhead I, Herrero J, Hobolth A, Lappalainen T, Mailund T, Marques-Bonet T, et al. 2012. Insights into hominid evolution from the gorilla genome sequence. Nature 483:169–175.

Tak PP, Firestein GS, Tak PP, Firestein GS. 2001. NF-κB: a key role in inflammatory diseases. J Clin Invest 107:7–11.

Thornton GK, Woods CG. 2009. Primary microcephaly: do all roads lead to Rome? Trends Genet 25:501–510.

Vallender EJ, Lahn BT. 2006. A primate-specific acceleration in the evolution of the caspase-dependent apoptosis pathway. Hum Mol Genet 15:3034–3040.

Vandepoele K, Andries V, Van Roy N, Staes K, Vandesompele J, Laureys G, De Smet E, Berx G, Speleman F, van Roy F. 2008. A constitutional translocation t(1;17)(p36.2;q11.2) in a neuroblastoma patient disrupts the human *NBPF1* and *ACCN1* genes. PLoS One 3(5), e2207.

Vandepoele K, Van Roy N, Staes K, Speleman F, Van Roy F. 2005. A novel gene family NBPF: Intricate structure generated by gene duplications during primate evolution. Mol Biol Evol 22:2265–2274.

Vandepoele K, Staes K, Andries V, van Roy F. 2010. Chibby interacts with NBPF1 and clusterin, two candidate tumor suppressors linked to neuroblastoma. Exp Cell Res 316:1225–1233.

Zhou F, Xing Y, Xu X, Yang Y, Zhang J, Ma Z, Wang J. 2013. NBPF is a potential DNA-binding transcription factor that is directly regulated by NF-κB. Int J Biochem Cell Biol 45:2479–2490.

